# Mesopredators display behaviourally plastic responses to dominant competitors when scavenging and communicating

**DOI:** 10.1101/2020.01.20.913335

**Authors:** Yiwei Wang, Maximilian L. Allen, Christopher C. Wilmers

## Abstract

Mesopredators face interspecific competition and intraguild predation when sharing resources with apex predators or more dominant mesopredators. We theorize that mesopredators use a variety of tactics to avoid competitively dominant predators at shared locations, such as scavenging and communication sites, that provide a mix of risks and rewards to these smaller predators. We examined whether mesopredator species employed behavioural tactics to reduce risks from dominant pumas when exploiting resources. We monitored carcasses in the Santa Cruz Mountains, CA across a gradient of human development and treated half of the carcasses with puma sign. Bobcats visited treated carcasses significantly later and for less time. Contrary to our expectations, coyotes and grey foxes were more likely to visit treated carcasses, although foxes were significantly less likely to visit a carcass also used by coyotes. Bobcats and foxes were less likely to visit carcasses at higher development levels whereas raccoons exhibited the opposite pattern. At communication sites, we observed temporal segregation among mesopredators and pumas. Coyotes and small predators exhibited the most segregation, followed by coyotes and pumas, and raccoons and pumas. Our results suggest subordinate predators employ a combination of spatial and temporal avoidance to minimize competitive interactions at shared sites.

---

Apex predators generally occur at low population densities but have disproportionate effects on their ecological communities (Estes 1996; Ripple et al. 2014). Predators affect prey species directly through predation (Estes et al. 2011; Ripple et al. 2014) or indirectly by inducing behavioural changes, such as altering prey habitat use (Altendorf et al. 2001; Brown et al. 1999). Antagonistic interactions also occur among members of the predator guild (Allen et al. 2016a), and lethal or competitive controls amongst predators often take a hierarchical form in which smaller predators are subordinate to larger ones (Estes 1996; Ripple et al. 2014). Intraguild cascades are therefore triggered by apex predators and influence both the populations and behaviours of subordinate predators. For example, Levi and Wilmers (2012) found that grey wolves (*Canis lupus*) decreased populations of coyotes (*Canis latrans*), which in turn increased populations of red foxes (*Vulpes vulpes*).

Mesopredators respond to threats of interspecific competition and intraguild predation from larger ones by adopting a variety of behavioural tactics, including spatial avoidance (Fedriani et al. 1999). However, certain locations, including communication sites (e.g., scent posts) and ephemeral food sources (e.g., carcasses), may be attractive to both mesopredators and apex predators (Allen et al. 2015a; Allen et al. 2017a; Li et al. 2013; Selva et al. 2005; Wilmers et al. 2003). Despite the risk of encountering larger predators, mesopredators predators regularly visit these areas, which suggests they gain advantages that outweigh the potential risks of intraguild competition and predation. During visits to communication sites predators likely acquire useful information conveyed by intra- and inter-specific scent marks while also leaving their own scent marks for mate attraction and territorial defence (Allen et al. 2017b; Begg et al. 2003; Smith et al. 1989, Krofel et al. 2017). At carcasses, individuals receive direct energetic gains by scavenging carrion (Allen et al. 2015b; Selva et al. 2005; Wilmers et al. 2003). Because several predator species potentially share these resources these areas are ideal for studying species-specific behavioural tactics employed to minimize risk from encountering dominant competitors.

Across much of North America, pumas are the apex predators in a guild that includes several mesopredators. Pumas affect the populations and behaviours of other species in their ecological communities, through both killing their competitors (Ruth and Murphy 2009), but also provisioning mesopredators with carrion (Allen et al. 2015b). Pumas are 2-5 times bigger than the next largest mesopredators, which falls within the size differential range with the highest rates and intensities of interspecific killings (Donadio and Buskirk 2006). Among mesopredators, raccoons (*Procyon lotor*), grey foxes (*Urocyon cinereoargenteus*), Virginia opossums (*Didelphis virginiana*) and skunk species, have a similar subordinate relationship to coyotes and bobcats (*Lynx rufus*). In general, we expected subordinate mesopredators to employ behavioural tactics to mitigate dangers when forced to share resources with dominant predators. To test this theory, we examined how mesopredators in the Santa Cruz Mountains (Figure 1) responded to puma signs and cues, as well as to each other, by monitoring the spatiotemporal patterns of resource use for sympatric predators engaging in two types of behaviours: scavenging and communicating.

**Figure 1.**
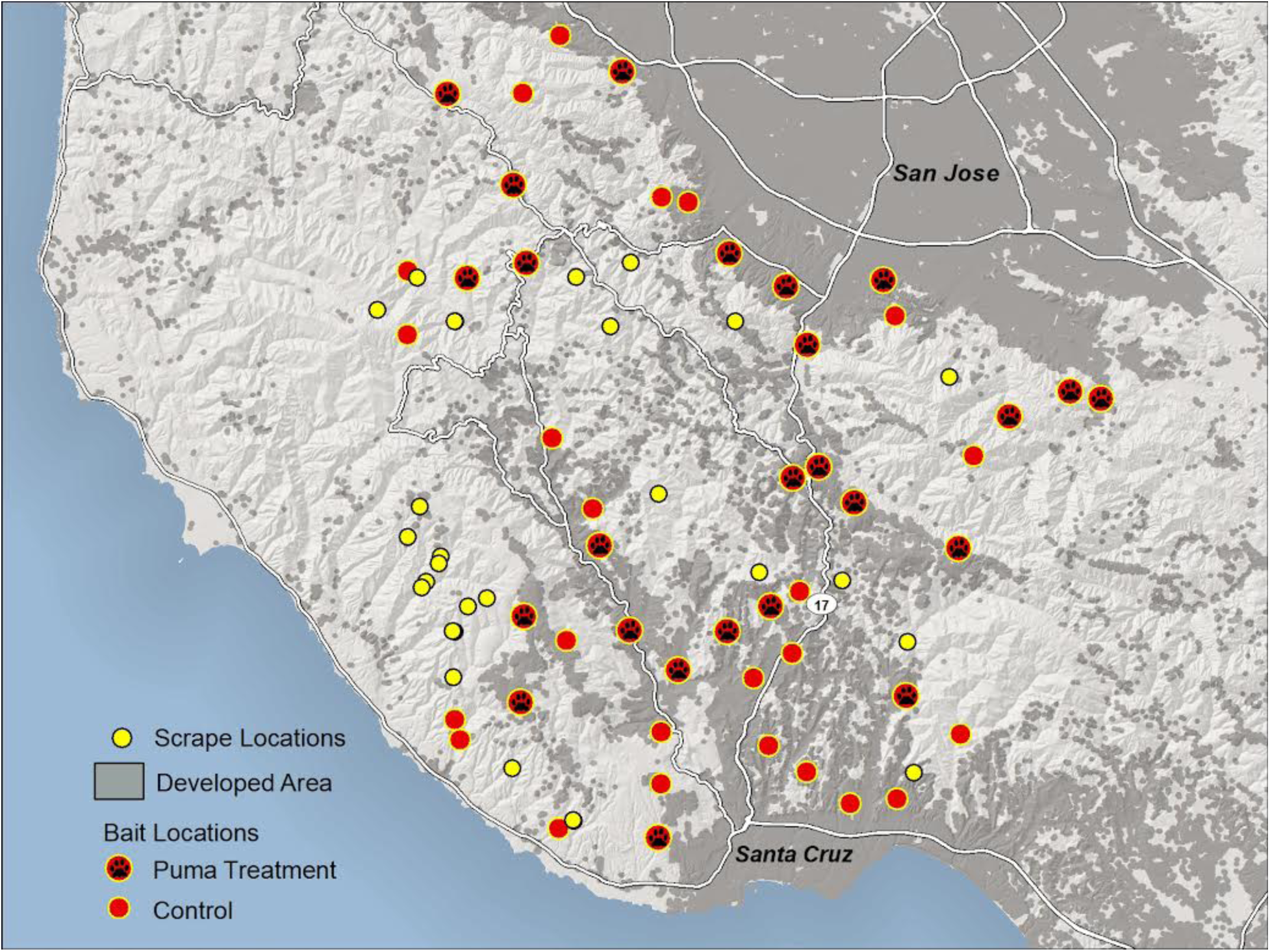
A map of the study area, which included areas in Santa Cruz, San Mateo, and Santa Clara Counties in California. Developed areas are shaded as grey, and the location of each carcass and community scrape area is noted.

To test for avoidance behaviour while scavenging, we used a field experiment to observe how mesopredators responded to simulated puma sign at black-tailed deer (*Odocoileus hemionus columbianus*) carcasses distributed across a gradient of human development. Carrion locations are likely sites for interspecific interactions among carnivores, as species compete to access a temporary and limited resource. While mesopredators can gain substantial energetic benefits from carrion, they may employ species-specific strategies to avoid pumas while accessing these resources (e.g., Allen et al. 2015b). We predicted that the mesopredators would generally avoid visiting or reduce their use of carcasses that showed signs of also being utilized by pumas or other dominant competitors. However, in developed areas with lower levels of puma activity, we expected that this avoidance behaviour would decline because carnivores display high behavioural plasticity and there would be fewer negative interspecific interactions with pumas (Crooks 2002; Wang et al. 2015; Wilmers et al. 2013), and they may be less fearful of pumas if they have reduced opportunities to interact with them.

We deployed motion-triggered cameras from 2010-2013 to document visits by pumas and mesopredators to community scrapes to collect evidence of avoidance behaviour. Community scrapes are areas regularly used for scent marking by pumas (Allen et al. 2014; Logan and Sweanor 2001) and the rest of the predator community (Allen et al. 2015a, Allen et al. 2017a). Many predator species predominately communicate by scent marking due to their relatively low levels of opportunities for direct intraspecific interactions (Bekoff et al. 1984; Logan and Sweanor 2001; Smith et al. 1989; Taylor et al. 2015). However, communication comes with risk, as signals may attract unwanted attention from more dominant predators and increase predation risk to the animal engaging in scent marking (Hughes et al. 2012; Hughes et al. 2010; Moller et al. 2011). As such, we hypothesized that mesopredators would display different activity patterns during their visits to scent communication sites with larger predators in order to reduce potentially lethal encounters. The preponderance of puma scent communication sites, or “community scrapes”, are located in relatively undeveloped habitats (Wilmers et al. 2013). As such we did not examine the degree to which these behaviours covaried with development at these sites.

## Methods

### Study area

We conducted our study in a 1,700 km^2^ study area in the Santa Cruz Mountains of California (Figure 1). The study area is bounded by the Pacific Ocean to the west and Highway 101 to the east, with the cities of San Francisco and San Jose to the north. The study area is divided by a major highway (California Highway 17), which was an important source of mortality for pumas (Wilmers et al. 2013) and other predators. Elevation ranged from sea level to 1,155m, and the climate is a mild Mediterranean. Historical average daily high temperatures ranged from 15.5–24.4°C and average daily low temperatures ranged from 3.9–11.1°C. Most of the rainfall occurs between November and April, resulting in average of 58-121 cm (Wilmers et al. 2013). Other predators in the study area include bobcats, coyotes, grey foxes, raccoons, Virginia opossums, spotted skunks (*Spilogale gracilis*), and striped skunks (*Mephitis mephitis*).

### Compliance with Ethical Standards

All field experimental procedures involving live animals were carried out in accordance with approved guidelines from the Independent Animal Care and Use Committee at the University of California, Santa Cruz (Protocols Wilmc0709 and Wilmc1101). All protocols were performed within the guidelines set by the University of California and the American Society of Mammalogists. The authors have no known conflicts of interest.

### Scavenging experiment design and analyses

We implemented a field experiment with motion-triggered cameras (Bushnell TrophyCam, Overland Park, KS) to evaluate whether mesopredators responded to signs of pumas while scavenging at black-tailed deer carcasses in a human-altered landscape. We used hind legs (weighing 4.5-7.0 kg) from road-killed deer as carcasses. We deployed 49 carcasses (Figure 1) in the study area between February and April 2012, and used thick rope to secure carcasses and prevent mesopredators from immediately dragging them away. We placed the carcasses randomly along a stratified development gradient of approximately 0-2 houses per hectare, but at a distance of at least 1km apart. Housing density was determined by manually digitizing houses from satellite images or using a street address data layer for urban areas in ArcGIS (v. 10.0, ESRI 2010; see Wang et al. 2015). To simulate the presence of pumas at carcasses, we treated 25 of 49 carcasses with puma faeces (placed 1m away from the carcass) and 4 mLs of puma urine, distributed evenly at orthogonal angles on all four sides of the carcass within a 1m radius. We obtained the faeces from a captive puma in a nearby sanctuary (Wild Things, Salinas, CA) and urine from In Heat Scents (Kinston, AL, USA). The faeces were stored in a freezer, then defrosted and placed at carcass sites within a week of collection, and the urine was stored in a standard refrigerator. Each carcass was monitored by a camera, which was set to capture one picture at a time with a minimum 1 sec delay between photos. We monitored each carcass for 14 days, and then removed the cameras.

We recorded all predator species that visited each carcass, the time of the first visit by each species, the time of day for the start of each visit, and the duration of each visit. We used generalized linear models (GLMs) in the program R (v3.0.0, R Core Team 2013) to compare how puma treatment, housing development, and local forest cover affected whether mesopredator species visited carcasses. We used a log link for a binomial error distribution, as puma treatment was coded as a binary variable, with 1 representing treatment and 0 control. For mesopredator species that were not coyotes, we included coyote presence as a binary predictor variable because coyotes are also important intraguild predators of smaller predators and can exert a strong influence on their behaviour (Wang et al. 2015; Wilson et al. 2010). We defined local forest cover as the percentage of forested habitat in a circular area of radius 100m around the site using California GAP vegetation data (Lennartz et al. 2008). We computed the number of houses at three scales surrounding the carcass site (radius *=* 100m, 500m, and 1000m). We first used a top-down model selection approach by including a full model with all predictor variables and interactions between puma treatment and housing and coyotes and housing. We used backwards elimination to remove any predictors or interaction terms with a significance level of *P* > 0.2 and used likelihood ratio tests to determine whether the reduced models significantly improved fit. Finally, we identified the housing density scale that best fit the data by finding the combination of covariates and housing density radius that minimized the Akaike Information Criterion (AIC) weight for the model (Anderson and Burnham 2002).

We also evaluated whether behaviours associated with carcass visitations were impacted by either puma treatment, forest cover, or housing density at all three radius values. We used multiple linear regression in program R and log-transformed response variables when necessary to fulfil assumptions of normality. We tested whether the covariates influenced three behaviours: latency to first detection, the cumulative time spent at the carcass over the two-week period, and the mean time spent per visit. As with our previous analyses, we used backwards elimination to remove variables with no explanatory power and selected the final model by minimizing AIC weight.

### Communication area design and analyses

We identified community scrapes visually (Allen et al. 2014) and documented visits to community scrapes by pumas and mesopredators from December 2010 to November 2013. We placed motion-triggered cameras with infrared flash (Bushnell TrophyCam, Overland Park, KS) at 28 community scrapes (Figure 1), all of which were >1 km from each other. We programmed cameras to record a 60 sec video when triggered with a 1 sec delay before becoming active again. We located community scrapes using a modification of a custom program for identifying kill sites, using locations acquired from GPS-collared male pumas (see Wilmers et al. 2013), and also opportunistically located community scrapes while performing other field work (Allen et al. 2014).

For each video, we recorded the date, time, and species present at the community scrape. To improve independence of the activity data, we removed visits from the analysis if they occurred within 30 minutes of a previous visit of the same species at a given camera location. Using the remaining photos, we followed kernel density estimation methods described by Ridout and Linkie (2009) to construct probability density distributions of activity levels for pumas, coyotes, bobcats, grey foxes, raccoons, striped skunks, and opossums. We then used these distributions to compare the amount of temporal overlap (Δ), which ranges from 0 for no overlap to 1 for complete overlap, between different species pairs. We used program R (v3.0.0, R Core Team 2013) and the *Overlap* package (Meredith and Ridout 2014) for these statistical analyses.

Because grey foxes, striped skunks, and opossums had very similar activity levels (overlap values between 0.86-0.95), we combined them into one group that we called “small carnivores”. We calculated overlap values between pumas and mesopredators (coyotes, bobcats, raccoons, and small carnivores). We then calculated overlap values between coyotes and small carnivores and bobcats and small carnivores. For pairs including coyotes and raccoons, we used the Δ_1_ estimator for small samples, and for all other pairs we used the Δ_4_ estimator as recommended by Ridout and Linkie (2009). The Δ_1_ and Δ_4_ differ in that they use different sets of integers to predict the overlap values, and simulation studies have shown that Δ_1_ produce less biased values for smaller sample sizes (Ridout and Linkie 2009). We calculated 95% confidence intervals for all overlap estimates from 500 bootstrapped samples.

## Results

We documented 564 visits by mesopredators to deer carcasses, resulting in 28,719 photos. A total of 36 unique species visited the carcasses, including bobcats (n=24), coyotes (n=22), grey foxes (n=13), striped skunks (n=13), opossums (n=11), dogs (n=9), raccoons (n=7), and domestic cats (n=5). Pumas also visited four carcasses.

We documented 824 visits by pumas and 1424 visits by mesopredators to community scrapes. We recorded 7 subordinate predator species visiting community scrapes, including grey foxes (n=545 total visits), striped skunks (n=360), bobcats (n=329), opossums (n=154), coyotes (n=20), raccoons (n=15), and spotted skunks (n=1).

### Scavenging experiment: Presence at the carcass

Carcasses with puma treatment were more likely to be visited by coyotes (ß*=*2.21, *p*=0.023), grey foxes (ß*=*3.25, *p*=0.041), and striped skunks, to a lesser degree (ß=2.08; p=0.073), were more likely to visit carcasses with puma treatment, whereas the presence of bobcats, raccoons, or opossums at carcasses were unaffected by puma treatment (Table 1). The interaction between puma treatment and housing density (at 500m) also influenced the presence of coyotes (ß*=*-0.022, *p*=0.039) at carcasses and marginally influenced that of striped skunks (Table 1). Generally, coyotes and striped skunks were more likely to visit carcasses with puma treatment when they were located in areas with higher housing densities, but less likely to visit control carcasses located in areas with higher housing densities. Raccoons were more likely to visit carcasses located in higher housing locations (ß*=*0.018, *p*=0.008 at 500m), whereas grey fox presence was negatively correlated with housing density (ß*=*-0.019, *p*=0.008 at 1000m).

**Table 1.**
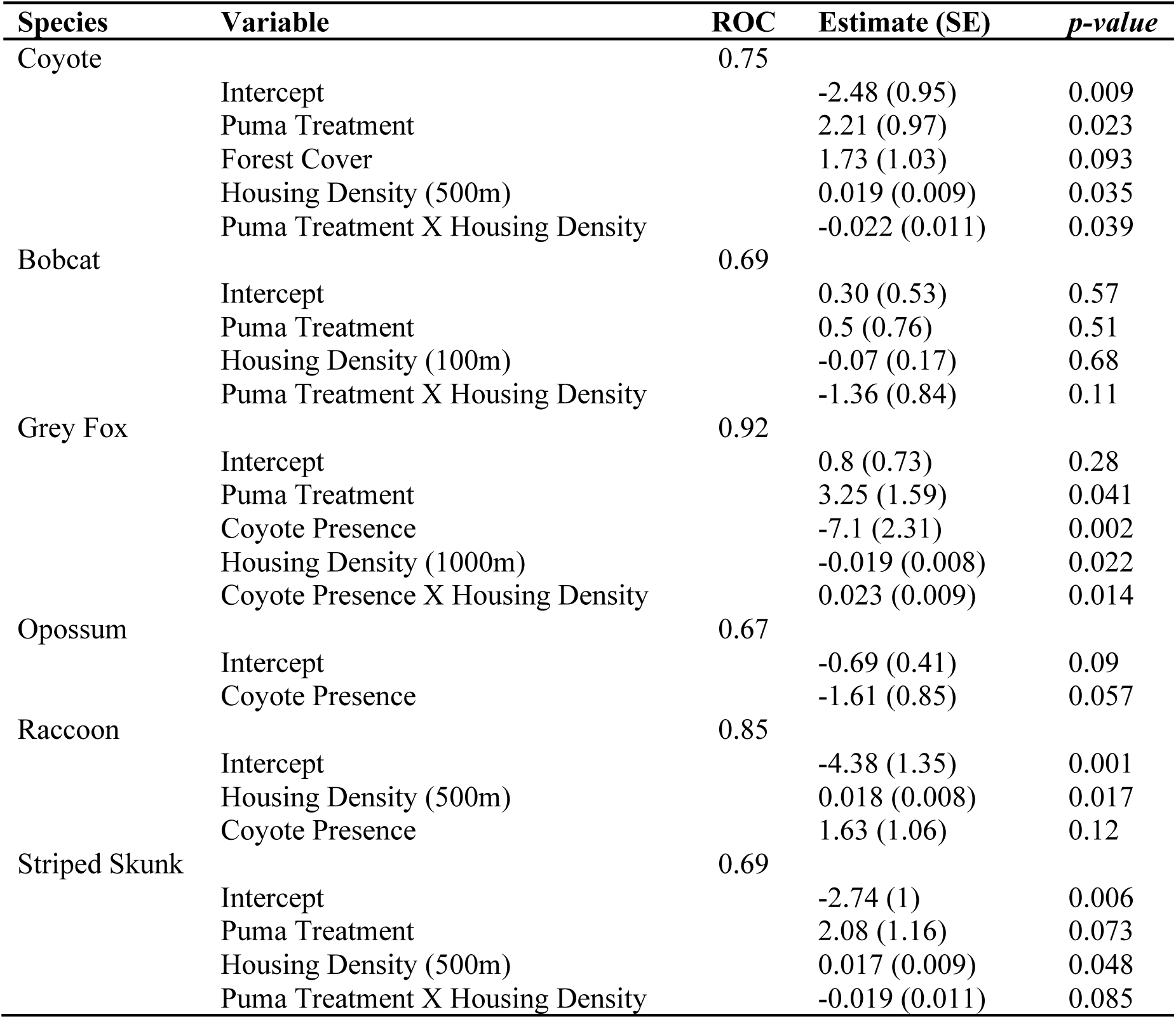
Logistic regression models for the probability of a species visiting carcasses. We tested the effects of puma treatment, forest cover, housing density, and coyote presence on the likelihood of each mesopredator species visiting carcasses. We report the receiver operating characteristic (ROC) for each of the best-fit models, along with the coefficient and its associated *p*-value for each variable.

Coyote presence was strongly negatively correlated with visitation by grey foxes (ß*=*-7.1, *p*=0.002), and was marginally negatively correlated with that of opossums. The best model explaining bobcat presence at carcasses had no significant terms, but included housing density (at 100m) and puma treatment (Table 1). While none of the covariates significantly correlated with bobcat presence, model predictions show bobcat presence declining with higher housing density.

### Scavenging experiment: Behaviour at the carcass

Bobcats were the only species that exhibited different behaviours at the carcass site in response to puma treatment; their latency to first detection was later (R^2^=0.22, *p*=0.013) and they spent less total time (R^2^=0.31, *p*=0.003) and had shorter visit durations (R^2^=0.28, *p*=0.004) at treated carcasses (Table 2). Raccoons visited carcasses earlier with both increased forest cover and housing density within a 100m radius (R^2^=0.80, *p*=0.018) (Table 2). Striped skunks increased the total amount of time they spent at carcasses with higher levels of housing (R^2^=0.32, *p*=0.025). Opossums spent more total time at carcasses with increased forest cover and housing density within a 100m radius (R^2^=0.51, *p*=0.024), and increased the mean duration of their visits at carcasses with puma treatment and higher forest cover (R^2^=0.51, *p*=0.024). Grey fox and coyote latency to detection and time spent at the kill were not significantly correlated with any of the covariates.

**Table 2.**
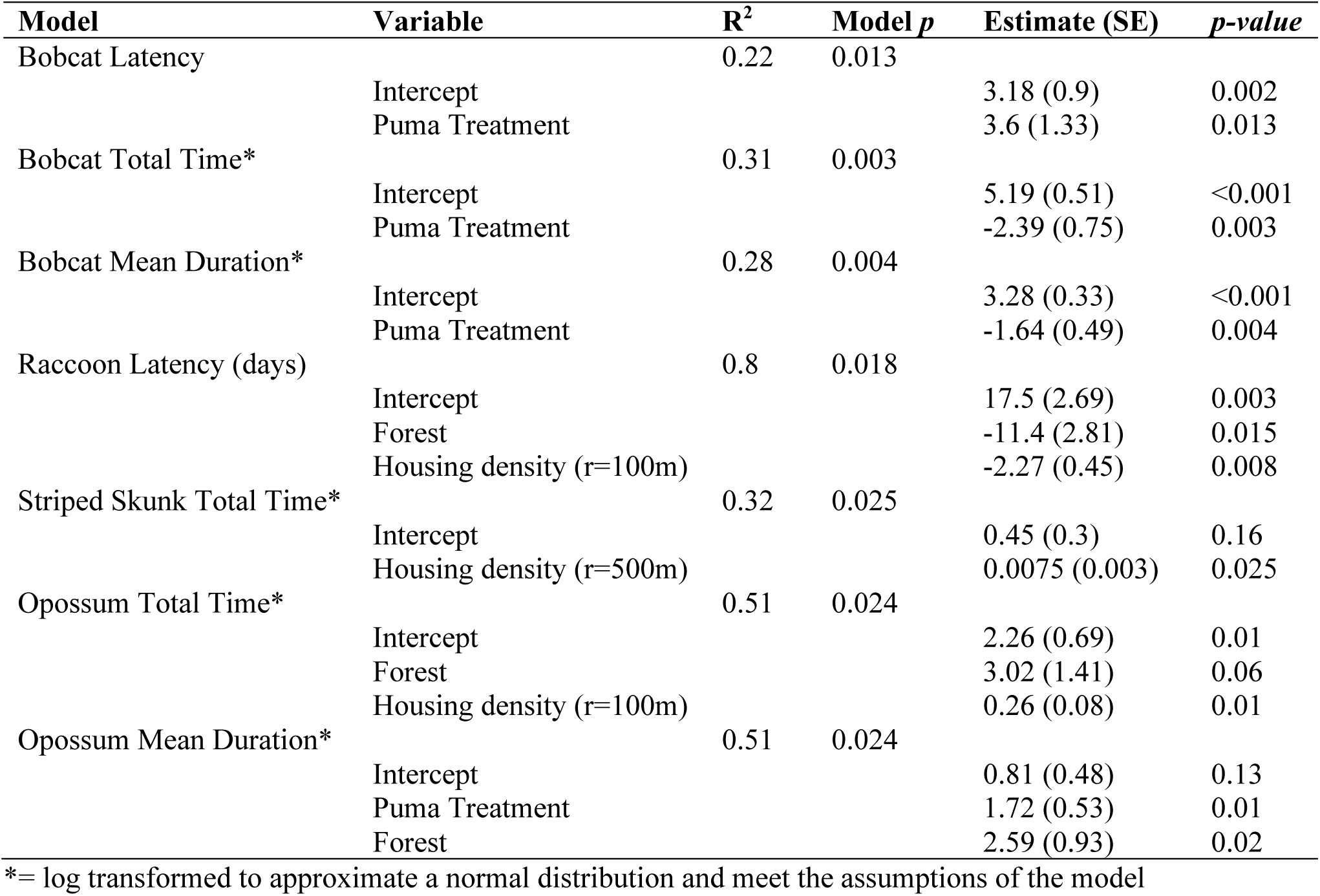
Best-fit regression models for mesopredator behaviours at our carcass experiments. We tested the effects of puma treatment, forest cover, and housing density on subordinate predator behaviour (i.e. latency to detection, total time at carcass, and mean duration of visit) at carcasses. We report the best model for each species and behaviour, in which the behaviour was significantly correlated with one or more covariates, and the coefficient and *p*-value for each variable.

| Model | Variable | R<sup>2</sup> | Model *p* | Estimate (SE) | *p*-value |
| --- | --- | --- | --- | --- | --- |
| Bobcat Latency | Intercept | 0.22 | 0.013 | 3.18 (0.9) | 0.002 |
|  | Puma Treatment |  |  | 3.6 (1.33) | 0.013 |
| Bobcat Total Time* | Intercept | 0.31 | 0.003 | 5.19 (0.51) | <0.001 |
|  | Puma Treatment |  |  | -2.39 (0.75) | 0.003 |
| Bobcat Mean Duration* | Intercept | 0.28 | 0.004 | 3.28 (0.33) | <0.001 |
|  | Puma Treatment |  |  | -1.64 (0.49) | 0.004 |
| Raccoon Latency (days) | Intercept | 0.8 | 0.018 | 17.5 (2.69) | 0.003 |
|  | Forest |  |  | -11.4 (2.81) | 0.015 |
|  | Housing density (r=100m) |  |  | -2.27 (0.45) | 0.008 |
| Striped Skunk Total Time* | Intercept | 0.32 | 0.025 | 0.45 (0.3) | 0.16 |
|  | Housing density (r=500m) |  |  | 0.0075 (0.003) | 0.025 |
| Opossum Total Time* | Intercept | 0.51 | 0.024 | 2.26 (0.69) | 0.01 |
|  | Forest |  |  | 3.02 (1.41) | 0.06 |
|  | Housing density (r=100m) |  |  | 0.26 (0.08) | 0.01 |
| Opossum Mean Duration* | Intercept | 0.51 | 0.024 | 0.81 (0.48) | 0.13 |
|  | Puma Treatment |  |  | 1.72 (0.53) | 0.01 |
|  | Forest |  |  | 2.59 (0.93) | 0.02 |

\*= log transformed to approximate a normal distribution and meet the assumptions of the model

### Communication site activity patterns

Kernel density estimates for predator species showed temporal segregation among several species pairs (Figure 2). Puma visits to community scrapes peaked during crepuscular hours and were primarily nocturnal (Figure 2). Bobcats and coyotes showed the most cathemeral patterns of all carnivore species, while smaller mesopredators (grey foxes, striped skunks, and opossums) visited scrapes almost exclusively during nocturnal hours (Figure 2). Raccoons were rare visitors to scrapes and were primarily observed nocturnally, but they rarely visited during the twilight hours when pumas were most active.

**Figure 2.**
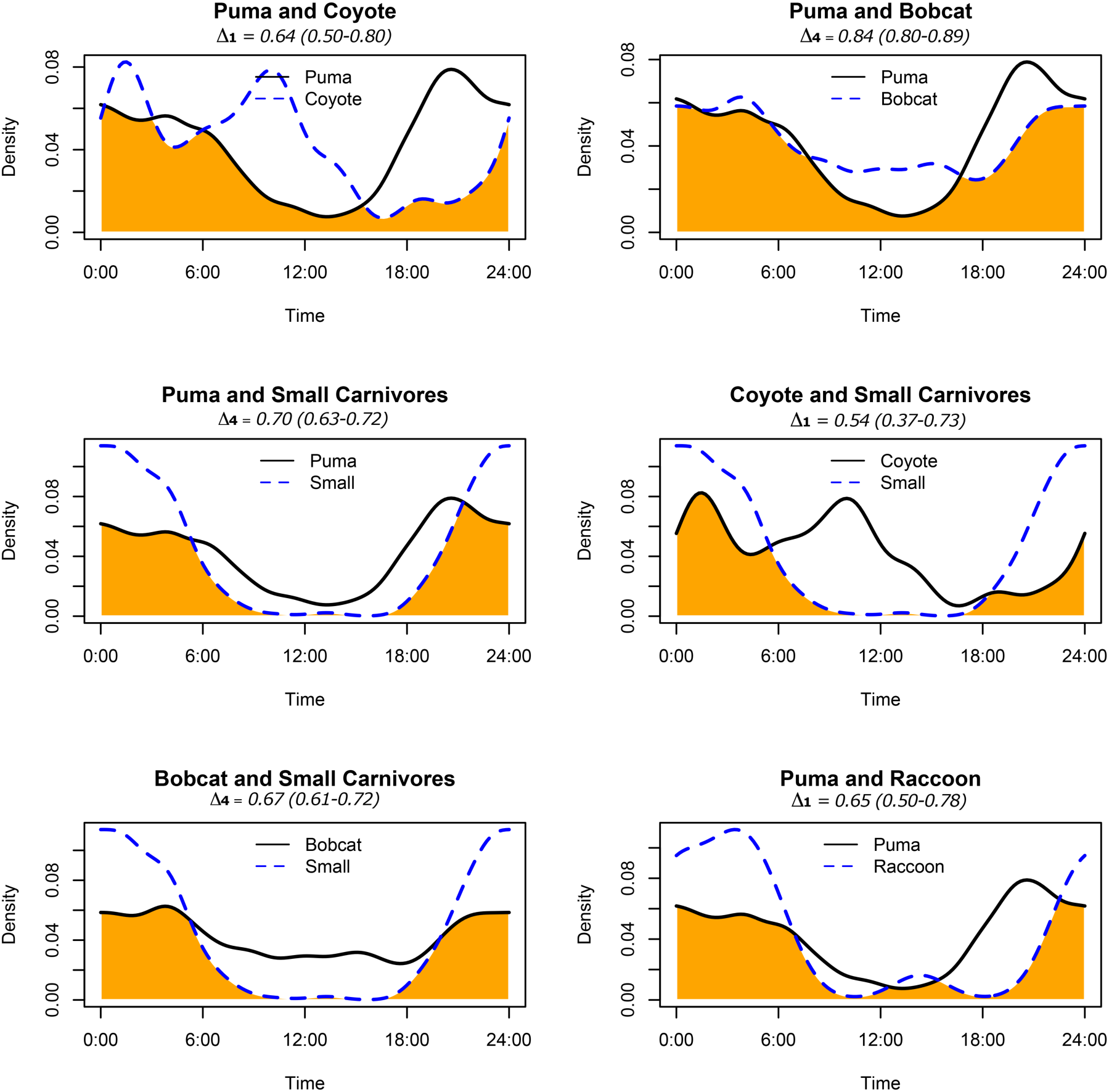
Temporal overlap (in orange) between species visiting community scrapes across a 24-hour period. Temporal overlap values (Δ) are listed along with 95% confidence intervals within parenthesis.

Aside from the similar temporal activity space shared by small carnivore species, bobcats and pumas exhibited the most temporal overlap of all species pairs at scrape sites (Δ_4_=0.84) indicating that bobcats were not avoiding times when pumas were most active at scrapes. Coyotes and raccoons were the least frequent visitors to community scrapes, and also had the lowest temporal overlap with pumas (coyote Δ_1_=0.64, raccoon Δ_1_=0.65). Of all species pairs, small carnivores exhibited the least amount of overlap with coyotes (Δ_1_= 0.54) and higher overlap with bobcats (Δ_4_=0.67) and pumas (Δ_4_=0.67).

## Discussion

Apex predators have strong effects on ecological communities, and evidence is emerging that their presence sometimes benefits smaller predators and prey species by suppressing mesopredator activity and populations (Levi and Wilmers 2012; Newsome and Ripple 2015). In a complex system containing multiple predator species, relationships between species pairs are complex and rarely linear. Our results highlight the behavioural diversity of mammalian predators in their responses to intraguild predation and competition and the strategies they employ to coexist with dominant predators at a fine spatial scale. Mesopredator species place themselves at considerable risk by visiting or consuming resources that may also attract apex or more dominant predators. In order to manage this risk, mesopredators can either avoid habitats and resources used by superior competitors or change their activity to avoid encountering these predators (Wang et al. 2015). Our study examined how mesopredators utilized carcasses and communication sites, locations that attract multiple predator species. While all mesopredator species were present throughout the study area (Wang et al. 2015), we found that mesopredators exhibited a variety of spatiotemporal avoidance tactics to potentially facilitate coexistence with pumas and other dominant predators. These tactics may vary depending on whether the individual was scavenging or communicating, since there are different risks and rewards associated with different behaviours. It is unclear why some species respond more strongly to competitive pressures than others, but some of this is likely due to the evaluation of the risks versus rewards of visiting specific resources.

Bobcats, coyotes, and grey foxes were the most common mammalian scavengers at our experimental carcasses, followed by opossums, striped skunks, and raccoons. Mesopredators visited carcass locations treated with puma sign at equal or higher probability than control carcasses, which was counter to our predictions. We expected that these small predators would avoid kill sites that had obvious puma cues, especially because pumas tend to return to their kills over a period of days (Allen et al. 2015c, Logan and Sweanor 2001). This may be due to an experimental flaw from not accounting for puma body odour at the sites, which has been shown to be an important source of information for mesopredators (Garvey et al. 2016; Leo et al. 2015). Alternatively, some species may have been attracted by puma faeces and urine because they associated them with an important food resource (e.g., Allen et al. 2015b). For example, Garvey et al. (2016) observed interspecific eavesdropping by stoats, which were attracted to food locations treated with odours from more dominant predators. However, we did find evidence of grey foxes and opossums avoiding carcasses utilized by coyotes, indicating that the rewards of scavenging do not always outweigh the risks, as coyotes can be a major source of grey fox mortality (Fedriani et al. 2000). Raccoons are frequent scavengers (DeVault et al. 2011; Olson et al. 2012), but they avoided carcasses treated with puma sign and their relative scarcity during these studies may hint at spatial avoidance of predator hotspots, which could be expected as raccoons are a common prey species in puma diets. Only bobcats exhibited a behavioural response to puma treatments by arriving later to feed at carcasses treated with puma sign and stayed for shorter periods on average and cumulatively. As the only other native felid in the ecosystem, it is possible that bobcats were more sensitive to puma sign than the other scavengers were, possibly supporting several studies have documented temporal partitioning among sympatric felid species (Di Bitetti et al. 2009; Harmsen et al. 2009).

We found evidence that development may have influenced whether predators visit carcasses because bobcat and grey fox presence at carcasses was negatively correlated to development levels. Surprisingly, bobcats were less sensitive to development (*h =* 100m) than grey foxes (*h =* 1000m), which is contrary to findings from previous studies (Bidlack 2007; Crooks 2002; Crooks and Soulé 1999). Raccoons, conversely, were more likely to visit carcasses in areas with higher housing. Interestingly, coyotes responded more positively to puma cues at higher housing densities, suggesting that their perception of risk may be altered in human dominated landscapes (Newsome et al. 2015). Pumas are less active in more developed areas (Wilmers et al. 2013), so it is possible that coyotes living in these areas have fewer competitive interactions with pumas and are less fearful of them, or are naïve of pumas and attracted to the novelty of their scent. Another possibility is that coyotes do not often encounter puma scents in developed areas and may thus be more attracted to the odour either as a novel stimulant or as a rare source of information on the presence or status of apex predators (Garvey et al. 2016).

Although we did not quantify spatial segregation (i.e., varied use of different scrape sites across the landscape), it appears that subordinate predators primarily used temporal segregation to avoid dominant competitors at community scrapes. While all predators showed a preference for nocturnal visits, temporal segregation among pumas, coyotes, and smaller predators was evident. The patterns of temporal segregation by pumas and mesopredators at community scrapes may suggest a cascading pattern initiated by pumas. Although our study area includes a mix of developed and undeveloped sites, there is still a high background level of human activity pervasive across the region due to its small size. Thus, we expected the predators in our study system to skew towards higher nocturnal activity. Pumas visited scrapes at all times of day, but were most active in the period after dusk, and generally throughout the night (Figure 3). In contrast, small predators appeared to be primarily strict nocturnal species whereas coyotes and bobcats were facultative cathemeral species (Figure 3) (Monterroso et al. 2014). While we don’t know whether coyotes increased their diurnal activity specifically to avoid pumas, the phenomenon of predators changing their activity to avoid encountering apex carnivores, such as humans, is well documented (Ordiz et al. 2012; Wang et al. 2015). Because community scrapes were located in areas with low human influence (e.g. Wilmers et al. 2013), we expect that the temporal pattern of puma activity reflected their natural preference, rather than one directed by humans.

**Figure 3.**
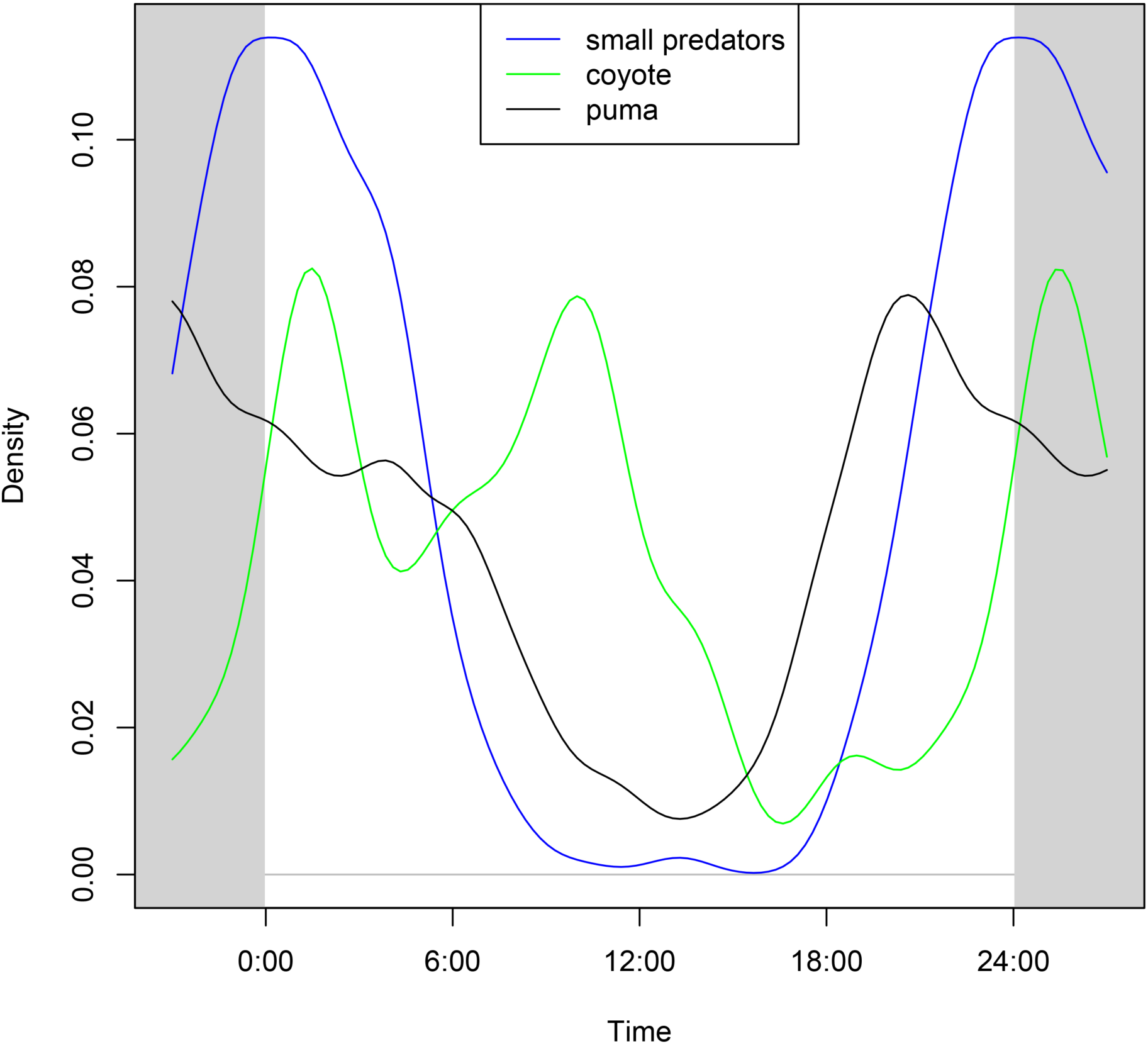
The proportion of visits to scrape sites at given times for pumas (apex predator), coyotes (mesopredator), and small predators. The figure illustrates a potential behavioural cascade, where pumas are primarily crepuscular, coyotes shift to become more cathemeral to potentially avoid pumas, and small predators are more strictly nocturnal to potentially avoid coyotes and pumas.

Because these diverse species occupy the same space and require similar resources, we expect that spatiotemporal partitioning may be one of the major behavioural responses to interspecific competition. Total avoidance of competitors is not possible, but we believe the temporal segregation practiced by the predator community partially facilitates coexistence. Small carnivores and coyotes exhibited the lowest amount of overlap, perhaps reflecting the competitive dominance of coyotes. Our findings support our hypothesis of mesopredator use of temporal segregation as a tool to facilitate coexistence. Some carnivores may just be using community scrapes for intraspecific communication, but other research shows that more complex interspecific communication is sometimes occurring (Allen et al. 2017a, Wilkenros et al. 2017). The shared use of community scrapes by carnivores, despite the apparent risks, raises the question of what benefits they receive from visiting those locations, and the effects of large carnivores on the communication behaviours of subordinate ones should also be considered in future studies.

## Acknowledgements

We thank the Mid-Peninsula Open Space District, California State Parks, Santa Cruz Land Trust, Santa Clara Parks, San Mateo Parks and many private landowners for giving us access to their land. Funding was provided by the American Society of Mammalogists, National Science Foundation grants #0963022 and *#*1255913, the Gordon and Betty Moore Foundation, and the Department of Environmental Studies at University of California Santa Cruz. We thank P. Houghtaling, Y. Shakeri, L. Hibbler and many undergraduate interns for their contributions to fieldwork, and B. Nickel and A. Cole for help obtaining GIS data.

